# Simple rapid *in vitro* screening method for SARS-CoV-2 anti-virals that identifies potential cytomorbidity-associated false positives

**DOI:** 10.1101/2020.10.13.338541

**Authors:** Kexin Yan, Daniel J. Rawle, Thuy T.T. Le, Andreas Suhrbier

**Affiliations:** QIMR Berghofer Medical Research Institute, Brisbane, Queensland. 4029, Australia; GVN Center of Excellence, Australian Infectious Disease Research Centre, Brisbane, Queensland 4029 and 4072, Australia

## Abstract

The international SARS-CoV-2 pandemic has resulted in an urgent need to identify new anti-viral drugs for treatment of COVID-19 patients. The initial step to identifying potential candidates usually involves *in vitro* screening. Here we describe a simple rapid bioassay for drug screening using Vero E6 cells and inhibition of cytopathic effects (CPE) measured using crystal violet staining. The assay clearly illustrated the anti-viral activity of remdesivir, a drug known to inhibit SARS-CoV-2 replication. A key refinement involves a simple growth assay to identify drug concentrations that cause cellular stress or “cytomorbidity”, as distinct from cytotoxicity or loss of viability. For instance, hydroxychloroquine shows anti-viral activity at concentrations that slow cell growth, arguing that its purported *in vitro* anti-viral activity arises from non-specific impairment of cellular activities.

---

The global SARS-CoV-2 pandemic has resulted in widespread activities seeking to identify new anti-viral drugs that might be used to treat COVID-19 patients^1–4^. Remdesivir has emerged as a lead candidate with clear anti-viral activity *in vitro*^5^ and non-human primates^6^, with some promising early results in human trials^7, 8^. The quest for new anti-viral drugs for SARS-CoV-2 (as for other viruses) usually begins with *in vitro* screening to identify potential candidates^9–11^. Initial screening usually involves assessing whether drugs can inhibit virus replication in a permissive cell line, with Vero E6 cells widely used for SARS-CoV-2. Such *in vitro* screening approaches often identify drugs that work well *in vitro*, but ultimately fail to have anti-viral activity *in vivo*. For example, chloroquine/hydroxychloroquine inhibits SARS-CoV-2 replication *in vitro*^5, 12^, but the drug ultimately emerged to have no utility in COVID-19 patients^13–15^. Chloroquine/hydroxychloroquine has been shown to have *in vitro* antiviral activity, but no antiviral activity in humans for a number of viruses including Epstein Barr virus (infectious mononucleosis)^16^, dengue^17^ HIV^18^, chikungunya^19^, Ebola^20^ and influenza^21^.

Although there are multiple reasons why *in vitro* anti-viral activity often does not translate into *in vivo* utility, one reason for false positives from *in vitro* screening assays is the misapplication of the therapeutic index concept as it applies to tissue culture-based anti-viral drug discovery, where this index is generally referred to as the selectivity index. The concentration of a drug that inhibits virus replication is often compared to the concentration that kills the cells (cytotoxicity). The MTS assay is also often used as a cytotoxicity or viability assay, although it actually measures mitochondrial activity. Viral replication would clearly be inhibited in cells that are not viable; however, what is perhaps under-appreciated is that viable, but stressed or otherwise slightly poisoned or compromised cells, are also likely to have a reduced capacity to replicate virus. Cellular stress responses can take multiple forms, but a key outcome of most stress responses is inhibition of translation^22–25^. Translational inhibition is also a key anti-viral response, which is able to inhibit replication of many viruses^22, 25, 26^ including coronaviruses^27^. Thus a drug that has no specific anti-viral activity, but that is able to induce cellular stress, may therefore inhibit virus replication non-specifically and generate a potential false positive in the screening assay.

A key outcome of stress responses is usually to slow cell growth, allowing the cell to either recover, or if stress and/or damage is excessive, to induce cell death^28–30^. Cells that are slightly poisoned or otherwise compromised (without induction of stress responses) would likely also show reduced growth rates. Cell growth of Vero E6 cells can be very simply measured by seeding 400 cells per well in triplicate into a 96 well flat bottom plate and culturing with a range of drug concentrations for 4 days followed by crystal violet staining. The percentage of protein staining relative to a no-drug control is then calculated and provides a simple measure of the drug concentration that slows cell growth. Perhaps not surprisingly the drug concentrations that caused inhibition of cell growth were usually lower than the drug concentrations that caused cytotoxicity (Fig. 1, compare black circles with green squares). For some drugs the concentration differences for these two activities were ≥10 fold (Fig. 1, ribavirin, cycloheximide, oleuropein, didemnin B). Inhibition of cell growth is not really cytostasis, which generally means no growth, and not really cytotoxicity, which is generally viewed as cell death. The reason(s) for reduced cell growth induced by any given drug may not be clear, and may be related to stress responses or some other phenomena that compromises the cells normal metabolic activities. Hence we suggest the term “cytomorbidity” to infer a level of cytotoxicity insufficient to kill the cells or induce cytostasis, but sufficient to stress or compromise the cells, with a simple growth bioassay used to indicate cytomorbidity.

**Figure 1.**
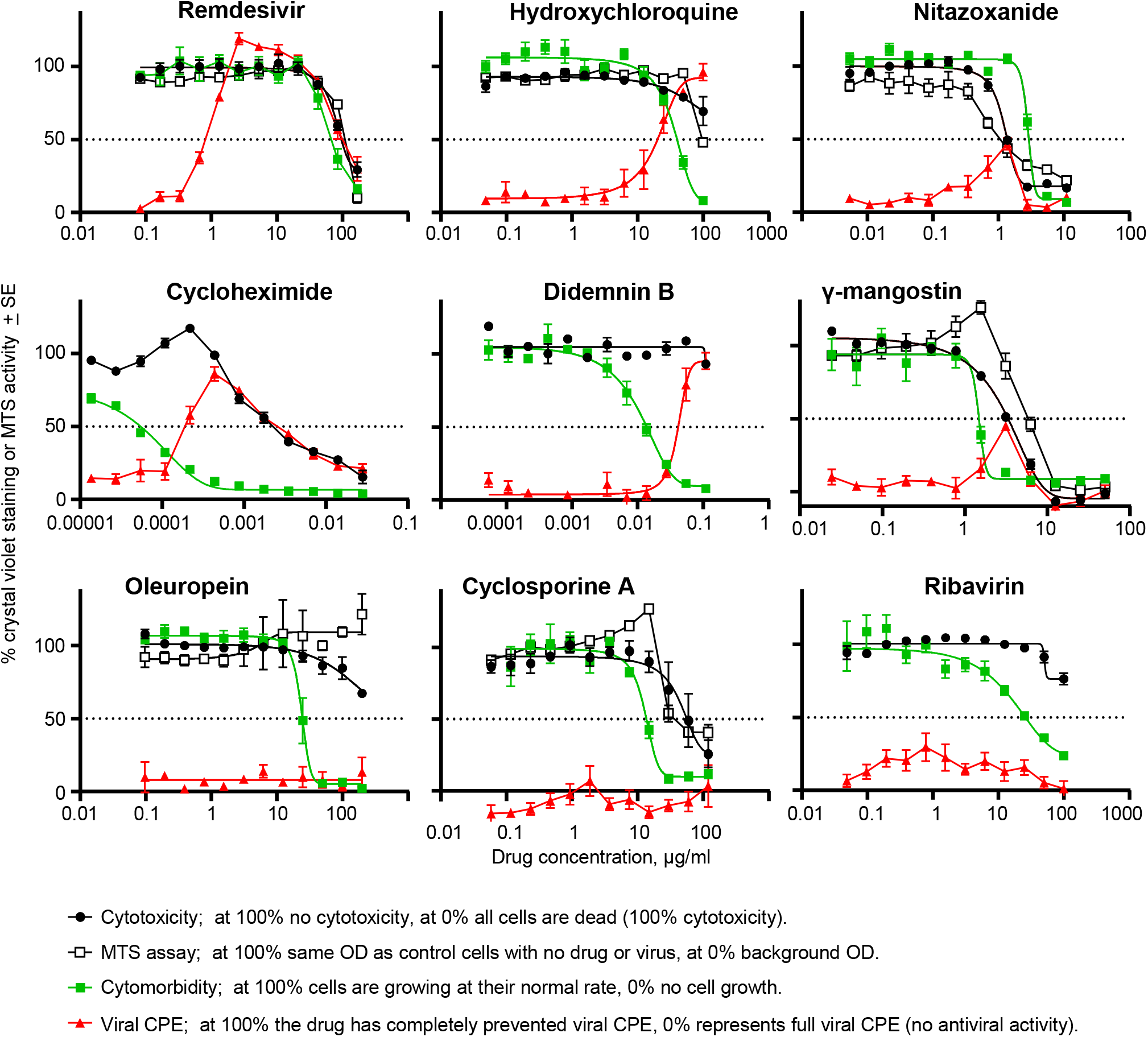
Drug cytotoxicity, cytomorbidity and inhibition of SARS-CoV-2-induced CPE. The indicated drugs at the indicated concentrations were cultured with Vero E6 cells (i) without virus and 10^4^ cells per well to measure cytotoxicity or MTS activity (black circles and white squares), (ii) without virus and 400 cells per well to measure cytomorbidity (green squares) or (iii) with virus and 10^4^ cells per well to measure viral CPE (red triangles). Error bars represent standard error of the mean (SEM) for 3-6 replicates, with each experiment undertaken independently in triplicate 1-2 times. The micromolar (μM) concentration of each drug at 1 μg/ml is; remdesivir = 1.66, hydroxychloroquine = 2.3, nitazoxanide = 3.25, cycloheximide = 3.56, didemnin B = 9.09, γ-mangostin = 2.52, oleuropein = 1.85, cyclosporine A = 0.83, ribavirin = 4.1.

A simple rapid bioassay for screening drugs for potential antiviral activity against SARS-CoV-2 is to determine whether the drug can inhibit virus-induced cytopathic effects (CPE) in Vero E6 cells. Remdesivir is known to inhibit SARS-CoV-2 replication^5^ and is used herein to illustrate the behavior of an effective drug in this bioassay. Remdesivir was able to inhibit virus-induced CPE by 50% at ≈1 μg/ml and the drug caused 50% cytotoxicity at ≈100 μg/ml, providing a selectivity index of ≈100. Importantly, remdesivir showed cytomorbidity at ≈80 μg/ml, which still leaves a selectivity index of ≈80 (Fig. 1 and 2, Remdesivir). Hydroxychloroquine was able to inhibit viral CPE by 50% at ≈20 μg/ml and showed a 50% loss of viability using the MTS assay at ≈100 μg/ml, suggesting a selectivity index of ≈5. However, cytomorbidity was clearly evident at ≈20 μg/ml, so the anti-viral activity occurred at similar concentrations to those that caused cytomorbidity (Fig. 1, Hydroxychloroquine); indicating a potential false positive. The overlapping activities are clearly evident when the crystal violet stained plates are viewed (Fig. 2).

**Figure 2.**
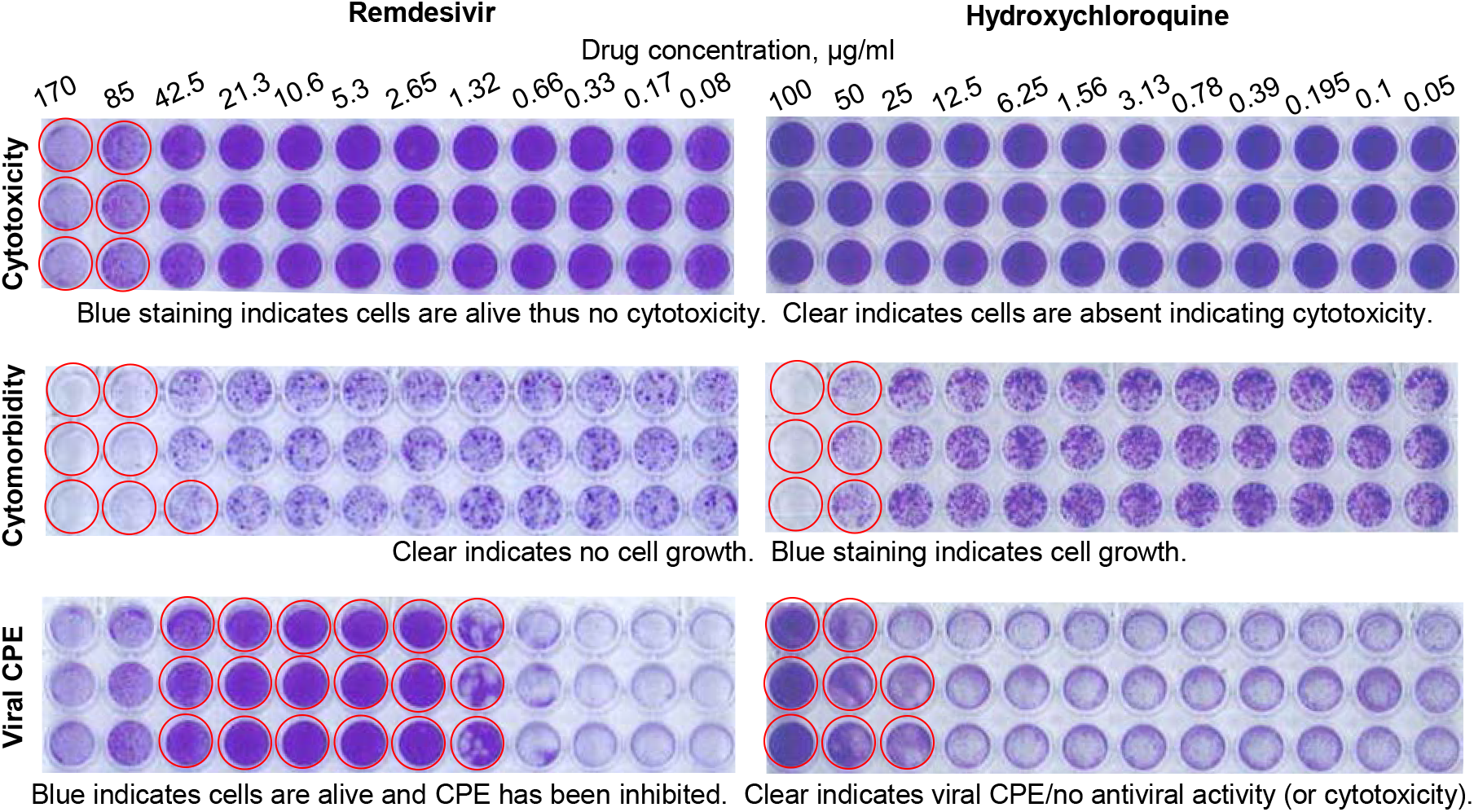
Crystal violet staining for remdesivir and hydroxychloroquine. Cytotoxicity assay (Vero E6 seeded at 10^4^ cells/well with no virus). Cytomorbidity assay (Vero E6 seeded at 400 cells/well with no virus). Viral CPE (Vero E6 seeded at 10^4^ cells/well with virus MOI≈0.01). After 4 days in culture 96 well plates were fixed and stained with paraformaldehyde and crystal violet respectively, washed in water, dried and scanned. For the Cytotoxicity assay wells encircled in red show overt cytotoxicity. For the Cytomorbidity assay wells encircled in red show overt cell growth reduction. For viral CPE assay, wells encircled in red show inhibition of CPE indicating potential antiviral activity.

The close relationship between anti-viral activity and translation inhibition (inherent in the stress responses described above) can be seen with the use of the translation inhibitors, cycloheximide and didemnin B. These drugs provide selectivity indices of ≥10, when comparing viral CPE inhibition and cytotoxicity. However, concentrations that inhibited viral CPE again overlapped with those that caused cytomorbidity (Fig. 1, Cycloheximide, Didemnin B). The drug γ-mangostin would appear to have a small level of anti-viral activity with a low selectivity index, but again this activity overlapped with the cytomorbidity (Fig. 1, γ-mangostin). Thus, as for hydroxychloroquine, the assay results for these latter drugs provide no supportive data for anti-viral activity, instead they suggest these drugs inhibit viral replication non-specifically by impairing cellular activities. Nitazoxanide showed some anti-viral activity, but this coincided with cytotoxicity, providing an example of the conventional cytotoxicity control that would be used to argue that the drug has no specific anti-viral activity and has a selectivity index of 1 (Fig. 1, Nitazoxanide). Curiously, higher concentrations of nitazoxanide were needed to inhibit cell growth than were needed to induce cytotoxicity; likely an example cell density associated toxicity.

The frequently used MTS assay, as expected, often gave results similar to those provided by the cytotoxicity assay. Importantly the MTS assay did not provide a measure of cytomorbidity, presumably because mitochondria largely remain active even in stressed cells and/or cells in G0 (cytostasis). For oleuropein, cyclosporine A and γ-mangostin, cytomorbidity was associated with an increase in MTS activity (Fig. 1). The MTS bioassay may thus provide slightly misleading information in this context; i.e. increased mitochondrial activity, rather than indicating increased cell numbers, can sometimes be associated with stress or mild toxicity.

The CPE based assay described herein is really only useful for screening drugs that target the virus directly. For instance, drugs whose mechanism of action requires induction of type I interferons, would be ineffective as Vero E6 cells do not make type I interferons. Another limitation of using virus-induced CPE as a read-out for anti-viral drugs is sensitivity. Higher drug concentrations are likely needed to prevent viral CPE (overwhelming infection resulting in cell death) than would be needed to inhibit viral replication as measured (for instance) by qRT-PCR of virus released into culture supernatants^31^. Nevertheless, the CPE-based assay represents a screening tool able rapidly to identify promising anti-viral candidates. More sensitive assays could be also envisaged for assessing cytomorbidity, such as measuring activation of stress factors such as ATF3^32^, analyzing cell cycle perturbations by flow cytometry or cell growth kinetics using the IncuCyte live-cell analysis system. However, the simple growth assay proposed herein allows rapid identification of drug concentrations that disrupt cellular activities/functions, which are often sufficient to inhibit viral replication non-specifically. The cytomorbidity assay thereby flags potential false positives.

## Online content

### Methods

#### PC3/BSL3 facilities

All infectious SARS-CoV-2 work was conducted in a dedicated suite within the PC3/BSL3 facility at the QIMR Berghofer MRI (Australian Department of Agriculture, Water and the Environment certification Q2326 and Office of the Gene Technology Regulator certification 3445). Work was approved by the QIMR Berghofer MRI Biosafety Committee (P3600).

#### Cells and SARS-CoV-2 virus

Vero E6 cells (C1008, ECACC, Wiltshire, England; Sigma Aldridge, St. Louis, MO, USA) were cultured in medium comprising RPMI1640 (Gibco) supplemented with 10% fetal calf serum (FCS), penicillin (100 IU/ml)/streptomycin (100 μg/ml) (Gibo/Life Technologies) and L-glutamine (2 mM) (Life Technologies). Cells were routinely checked for mycoplasma (MycoAlert Mycoplasma Detection Kit MycoAlert, Lonza) and FCS was assayed for endotoxin contamination before purchase^1^. The SARS-CoV-2 virus was kindly provided by Queensland Health Forensic & Scientific Services, Queensland Department of Health, Brisbane, Australia. The virus (hCoV-19/Australia/QLD02/2020) was isolated from a patient and sequence deposited at GISAID (https://www.gisaid.org/; after registration and login, sequence can be downloaded from https://www.epicov.org/epi3/frontend#1707af). Virus stock was generated by infection of Vero E6 cells at multiplicity of infection (MOI)≈0.01, with supernatant collected after 3 days, cell debris removed by centrifugation at 3000 x g for 15 min at 4°C, and virus aliquoted and stored at −80 °C. Virus titers were determined using standard CCID_50_ assays (see below). The virus was determined to be mycoplasma free using co-culture with a non-permissive cell line (i.e. HeLa) and Hoechst staining as described^2^

#### Virus titration by CCID_50_ assay

Virus was titrated using a standard Cell Culture Infectivity Dose 50 (CCID_50_) assay. Vero E6 cells were plated into 96 well flat bottom plates at 2×10^4^ cells per well in 100 μl of medium (see above). The following day 100 μl of virus was added in 10 fold serial dilutions in RPMI 1640 supplemented with 2% FCS, and the plates cultured for 4 days at 37°C and 5% CO_2_. To inactivate virus and stain the cells, 50 μl of formaldehyde (15% w/v) and crystal violet (0.1% w/v) (Sigma-Aldrich) was added per well to the 200 μl of medium already present in each well. Plates were left overnight inside a closed biosafety cabinet with lids on (biosafety cabinet UV switched on for 20 mins). The plates (lids off) and inverted lids were then exposed to 7.6 kJ/m2 UV-C. The lids were replaced and the plates sprayed with 80% (v/v) ethanol; plates were labelled with Alcohol Resistant Cryogenic Permanent Markers (Science Marker). The plates are then deemed decontaminated and removed from the PC3/BSL3 facility. The plates were washed in tap water, dried (photographed or scanned) and 100 μl/well of 100% methanol added and OD read at 595 nm. The virus titer was determined by the method of Spearman and Karber (a convenient Excel CCID_50_ calculator is available at https://www.klinikum.uni-heidelberg.de/zentrum-fuer-infektiologie/molecular-virology/welcome/downloads).

#### Drugs

Didemnin B was kindly provided by the Natural Products Branch, NCI (NSC 325319, Bethesda, MD, USA). Cycloheximide, nitazoxanide, ribavirin, hydroxychloroquine sulfate, γ-mangostin and oleuropein were all purchased from Sigma Aldrich. Remdesivir was purchased from AdooQ BioScience. Cyclosporine A (Merck Millipore) was dissolved in 100% EtOh. Ribavirin and hydroxychloroquine sulfate was dissolved in Ultrapure Distilled Water (Life Technologies). All other drugs were dissolved in DMSO (Sigma Aldrich). All drugs were aliquoted and stored at −80°C.

#### Cytotoxicity testing and MTS assay

Vero E6 cells were plated as above, 10^4^/well in triplicate in 100 μl medium and cultured overnight. The drug was diluted in 2 fold serial dilutions in RPMI 1640 supplemented with 2% FCS, and 50 μl was then added per well (at 4 times the indicated final concentration). A further 50 μl RPMI 1640 supplemented with 2% FCS was then added per well. The plates were cultured for 4 days, after which they were fixed and stained with crystal violet as above. A MTS assay was performed where indicated (before fixation and crystal violet staining) using CellTiter 96 AQueous One Solution Cell Proliferation Assay (MTS) (Promega) as per manufacturer’s instructions.

#### Cytomorbidity testing

Vero E6 cells were plated at 400 cells per well in 100 μl medium. All other steps were performed as described above for cytotoxicity testing.

#### Anti-viral activity testing

Vero E6 cells were plated as above, 10^4^/well in triplicate in 100 μl medium and cultured overnight. The drug was added at 4 times the indicated final concentration in 50 μl RPMI 1640 supplemented with 2% FCS. SARS-CoV-2 virus at a MOI≈0.01 was then added in 50 μl RPMI 1640 supplemented with 2% FCS. After 4 days of culture the cells were fixed and stained with crystal violet as above.

## Data availability

All data is provided in the manuscript and accompanying online content.

## Acknowledgements

We thank Dr I Anraku for his assistance in managing the PC3 (BSL3) facility at QIMR Berghofer MRI. We thank Dr Alyssa Pyke and Mr Fredrick Moore (Queensland Health, Brisbane) for providing the SARS-CoV-2 virus. We thank Dr David Harrich for help with reagents. We thank Clive Berghofer and Lyn Brazil (and others) for their generous philanthropic donations to support SARS-CoV-2 research at QIMR Berghofer MRI. A.S. holds an Investigator grant from the National Health and Medical Research Council (NHMRC) of Australia (APP1173880).

## Author contribution

KY and TTL undertook the experiments. DJR, AS supervised the experiments, analysed the data and obtained funding. AS: wrote the manuscript with input from DJR.

## References

1. Santos, I.A., Grosche, V.R., Bergamini, F.R.G., Sabino-Silva, R. & Jardim, A.C.G. Antivirals Against Coronaviruses: Candidate Drugs for SARS-CoV-2 Treatment? Front Microbiol 11, 1818 (2020).

2. Elshabrawy, H.A. SARS-CoV-2: An Update on Potential Antivirals in Light of SARS-CoV Antiviral Drug Discoveries. Vaccines (Basel) 8, 335 (2020).

3. Pillaiyar, T., Wendt, L.L., Manickam, M. & Easwaran, M. The recent outbreaks of human coronaviruses: A medicinal chemistry perspective. Med Res Rev EPub (2020).

4. Teoh, S.L., Lim, Y.H., Lai, N.M. & Lee, S.W.H. Directly Acting Antivirals for COVID-19: Where Do We Stand? Front Microbiol 11, 1857 (2020).

5. Wang, M. et al. Remdesivir and chloroquine effectively inhibit the recently emerged novel coronavirus (2019-nCoV) in vitro. Cell Res 30, 269–271 (2020).

6. Williamson, B.N. et al. Clinical benefit of remdesivir in rhesus macaques infected with SARS-CoV-2. Nature 585, 273–276 (2020).

7. Wang, Y. et al. Remdesivir in adults with severe COVID-19: a randomised, double-blind, placebo-controlled, multicentre trial. Lancet 395, 1569–1578 (2020).

8. Beigel, J.H. et al. Remdesivir for the Treatment of Covid-19 - Preliminary Report. N Engl J Med, EPub (2020).

9. Pizzorno, A. et al. In vitro evaluation of antiviral activity of single and combined repurposable drugs against SARS-CoV-2. Antiviral Res 181, 104878 (2020).

10. Touret, F. et al. In vitro screening of a FDA approved chemical library reveals potential inhibitors of SARS-CoV-2 replication. Sci Rep 10, 13093 (2020).

11. Caly, L., Druce, J.D., Catton, M.G., Jans, D.A. & Wagstaff, K.M. The FDA-approved drug ivermectin inhibits the replication of SARS-CoV-2 in vitro. Antiviral Res 178, 104787 (2020).

12. Liu, J. et al. Hydroxychloroquine, a less toxic derivative of chloroquine, is effective in inhibiting SARS-CoV-2 infection in vitro. Cell Discov 6, 16 (2020).

13. Elavarasi, A. et al. Chloroquine and Hydroxychloroquine for the Treatment of COVID-19: a Systematic Review and Meta-analysis. J Gen Intern Med, E Pub (2020).

14. Khuroo, M.S. Chloroquine and hydroxychloroquine in coronavirus disease 2019 (COVID-19). Facts, fiction and the hype: a critical appraisal. Int J Antimicrob Agents 56, 106101 (2020).

15. FDA Revokes Emergency Use Authorization for Chloroquine and Hydroxychloroquine (June 15, 2020). https://www.fda.gov/media/138945/download.

16. Updike, S.J. & Eichman, P.L. Infectious mononucleosis treated with chloroquine. A double-blind study of 40 cases. Am J Med Sci 254, 69–70 (1967).

17. Tricou, V. et al. A randomized controlled trial of chloroquine for the treatment of dengue in Vietnamese adults. PLoS Negl Trop Dis 4, e785 (2010).

18. Rodrigo, C., Fernando, S.D. & Rajapakse, S. Clinical evidence for repurposing chloroquine and hydroxychloroquine as antiviral agents: a systematic review. Clin Microbiol Infect 26, 979–987 (2020).

19. Roques, P. et al. Paradoxical Effect of Chloroquine Treatment in Enhancing Chikungunya Virus Infection. Viruses 10, 268 (2018).

20. Dowall, S.D. et al. Chloroquine inhibited Ebola virus replication in vitro but failed to protect against infection and disease in the in vivo guinea pig model. J Gen Virol 96, 3484–3492 (2015).

21. Paton, N.I. et al. Chloroquine for influenza prevention: a randomised, double-blind, placebo controlled trial. Lancet Infect Dis 11, 677–683 (2011).

22. Houston, R., Sekine, S. & Sekine, Y. The coupling of translational control and stress responses. J Biochem 168, 93–102 (2020).

23. Girardin, S.E., Cuziol, C., Philpott, D.J. & Arnoult, D. The eIF2alpha kinase HRI in innate immunity, proteostasis and mitochondrial stress. FEBS J, Epub (2020).

24. Vind, A.C., Genzor, A.V. & Bekker-Jensen, S. Ribosomal stress-surveillance: three pathways is a magic number. Nucleic Acids Res, gkaa757 (2020).

25. Eiermann, N., Haneke, K., Sun, Z., Stoecklin, G. & Ruggieri, A. Dance with the Devil: Stress Granules and Signaling in Antiviral Responses. Viruses 12, E984 (2020).

26. Dalet, A. et al. Protein synthesis inhibition and GADD34 control IFN-beta heterogeneous expression in response to dsRNA. EMBO J 36, 761–782 (2017).

27. van den Worm, S.H. et al. Development and RNA-synthesizing activity of coronavirus replication structures in the absence of protein synthesis. J Virol 85, 5669–5673 (2011).

28. Fulda, S., Gorman, A.M., Hori, O. & Samali, A. Cellular stress responses: cell survival and cell death. Int J Cell Biol 2010, 214074 (2010).

29. Sionov, R.V. & Haupt, Y. The cellular response to p53: the decision between life and death. Oncogene 18, 6145–6157 (1999).

30. Pietenpol, J.A. & Stewart, Z.A. Cell cycle checkpoint signaling: cell cycle arrest versus apoptosis. Toxicology 181-182, 475–481 (2002).

31. Yao, X. et al. In Vitro Antiviral Activity and Projection of Optimized Dosing Design of Hydroxychloroquine for the Treatment of Severe Acute Respiratory Syndrome Coronavirus 2 (SARS-CoV-2). Clin Infect Dis 71, 732–739 (2020).

32. Ku, H.C. & Cheng, C.F. Master Regulator Activating Transcription Factor 3 (ATF3) in Metabolic Homeostasis and Cancer. Front Endocrinol (Lausanne) 11, 556 (2020).

## References

1 Johnson, B. J. et al. Heat shock protein 10 inhibits lipopolysaccharide-induced inflammatory mediator production. J Biol Chem 280, 4037–4047 (2005).

2 La Linn, M., Bellett, A. J., Parsons, P. G. & Suhrbier, A. Complete removal of mycoplasma from viral preparations using solvent extraction. J Virol Methods 52, 51–54 (1995).

